# Hyperplastic Growth, Not Hydrostatic Distension, in Endolymphatic Hydrops in Humans Challenges the Classic View of Meniere’s Disease

**DOI:** 10.1101/2025.03.11.642574

**Authors:** Corey Bryton, Diana M. Correa, David Bächinger, Valentin Fankhauser, Nicole C. Kamber, MengYu Zhu, Jennifer T. O’Malley, Tyler T. Hickman, Paula Robles-Bolivar, Steven D. Rauch, Andreas H. Eckhard

## Abstract

Meniere’s disease (MD), a degenerative inner ear disorder, is characterized by debilitating episodic vertigo “attacks” and hearing fluctuations, progressing to permanent sensory impairment. The prevailing dogma attributes these symptoms to an abnormal inner ear fluid buildup—known as endolymphatic hydrops (EH)—with concomitant rise of fluid pressure and repetitive microtrauma to sensory epithelia. However, this pressure-based mechanism lacks direct experimental evidence and fails to explain key clinical aspects of MD—exposing a critical gap in our disease understanding. To revisit the fundamental nature of EH, we performed 3D reconstructive, machine-learning-enhanced histological analyses and immunohistochemistry on human postmortem inner ear specimens. Contrary to the classic theory, EH-affected epithelia showed no signs of pressure-induced change. Instead, we observed an up to four-to seven-fold increase in epithelial cell number (hyperplasia) in both early and advanced EH stages. Quantification of the hyperplastic epithelial surface area, as well as immunohistochemical localization of key fluid homeostasis-associated proteins in the hyperplastic epithelium suggest this epithelial hyperplasia may actively compensate for cell loss in the endolymphatic sac, a key site of MD pathology. These findings challenge the conventional view of EH as solely a pathological pressure phenomenon, instead revealing an unexpected massive cellular expansion of these epithelia, consistent with a coordinated compensatory cellular response aimed at preserving inner ear fluid homeostasis and function in a compromised environment. This paradigm shift introduces dual beneficial and detrimental roles for EH, and suggests new therapeutic avenues for MD focused on promoting compensatory tissue repair while preventing maladaptive remodeling.

**Significance Statement:** For over a century, the leading explanation for Meniere’s disease—a debilitating inner ear disorder causing vertigo and hearing loss—has been a buildup of fluid and pressure in the inner ear, analogous to conditions like glaucoma. However, this long-held theory has never been directly proven, and treatments based on reducing this supposed pressure have shown limited success. Our research challenges this traditional view, revealing that the expansion of endolymphatic spaces is not primarily a fluid pressure problem, but secondary to a complex cellular response. This fundamental shift in understanding Meniere’s disease opens new avenues for developing effective therapies to prevent and treat hearing loss and vertigo attacks.

## Introduction

Meniere’s disease (MD) (1) is a degenerative inner ear disorder initially presenting with intense bouts of vertigo (spinning sensation), hearing loss, and tinnitus (ringing in the ear). As the disease progresses and the inner ear degenerates, patients eventually experience a permanent loss of both hearing and balance functions, profoundly impacting their quality of life (2). For over a century, the prevailing dogma has centered on endolymphatic hydrops (EH)—an abnormal accumulation and presumed elevated pressure of endolymph fluid within the diseased inner ear— as the root cause of symptoms and disease progression (3-6). This theory, rooted in analogies to fluid hypertension disorders like open-angle glaucoma and hydrocephalus (7, 8), has been reinforced by decades of postmortem histological (4, 9) and in vivo MRI (10, 11) findings of enlarged inner ear fluid spaces in MD patients, interpreted as primary “fluid overaccumulation”, causing repetitive fluid pressure surges, mechanically distending sensory epithelia to the point of histologically detectable ruptures (3, 12). These ruptures in turn are believed to disrupt transepithelial ionic gradients essential for sensory transduction, triggering acute symptom episodes and ultimately driving the progressive degenerative disease course (3, 12, 13). Although widely accepted, several lines of evidence have cast doubt on this fluid overpressure theory. First, no empirical data confirm fluid overpressure in MD or link it directly to symptom episodes (14-16). Second, a clear disconnect exists between EH and MD: EH is observed in other inner ear conditions (16-18) and even in inner ears deemed healthy (19), while some MD patients do not present with EH (20), arguing against a direct causal link. Finally, therapies effective for reducing fluid overpressure in glaucoma and hydrocephalus have failed to alleviate symptoms or halt disease progression in MD (21, 22). These inconsistencies challenge the existence of endolymph fluid overpressure as a causal factor in MD pathogenesis, highlighting a critical gap in our understanding of EH and the need for a reexamination of MD pathogenesis.

To address this gap, we hypothesized that EH arises from active cellular processes, possibly as a response to pathological events in the endolymphatic sac (ES) (23), a non-sensory part of the inner ear epithelium with presumed critical functions for inner ear fluid homeostasis. We analyzed archival inner ear tissue from individuals with MD and other EH-associated conditions (18), using manual and machine-learning (ML)-based histological methods to investigate epithelial integrity, cell patterning, and density, along with immunohistochemical analyses of proteins relevant to fluid-homeostasis in hydropic and non-hydropic epithelia. We also revisited historical specimens that formed the basis of the fluid overpressure theory, performing serial-section 3D reconstructions for a more comprehensive examination of epithelial cell organization. Our findings revealed previously unrecognized cellular features within hydropic epithelial domains, most notably a significant increase in epithelial cell numbers that fully accounts for the observed hydropic expansions. These findings provide strong support for a new “tissue damage-driven compensatory cell response” model of EH, with profound implications for understanding MD pathophysiology and guiding future therapy research.

## Results

### Presence of supernumerary cells in hydropic inner ear epithelia

Histological cross-sections of the cochlea (Fig. 1A) and saccule (Fig. 1B) from patients with MD displayed hallmark features of EH, including enlarged endolymph fluid spaces (red shading) with expansion of Reissner’s membrane (RM) and saccular membrane (SM) epithelia. Two key observations led us to question the traditional fluid-overpressure-driven model of EH. First, the hydropic epithelia frequently appeared tangled (Figs. 1C, 1C′), a configuration not easily explained by uniform passive distension from elevated fluid pressure. Second, we identified supernumerary cells arranged in villous extensions within the hydropic epithelia of some specimens (Figs. 1D, 1D′). While previous reports on cell density increase in hydropic epithelia have yielded conflicting results (24, 25), these atypical cellular arrangements led us to hypothesize that epithelial expansion in EH may result from active cell proliferation rather than passive mechanical distension (Fig. 1E).

**Figure 1.**
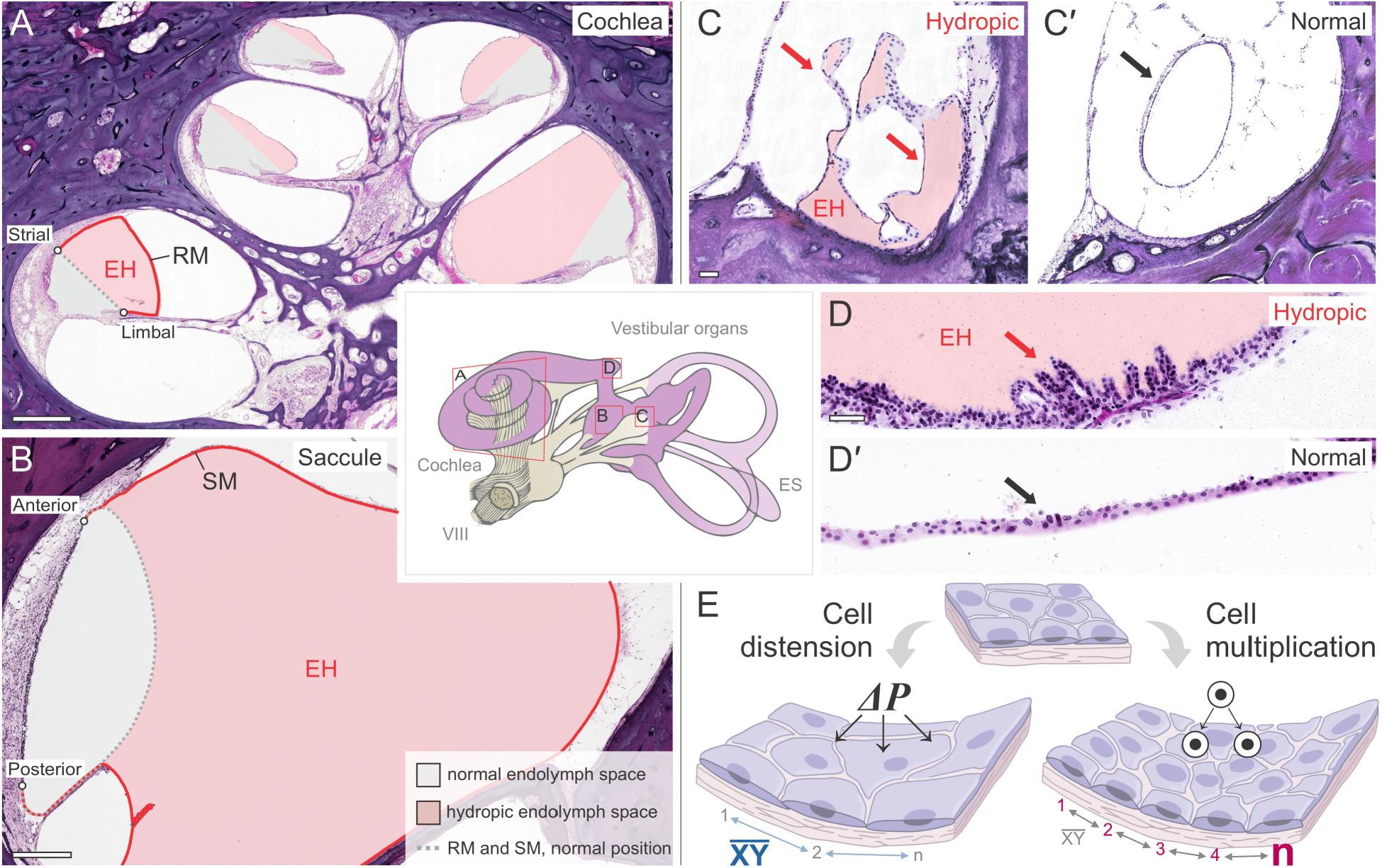
Histological features and contrasting models of epithelial expansion in endolymphatic hydrops (EH). (Center) Schematic of inner ear epithelium (purple), showing regions shown in histological cross-sections. (A, B) Cochlea (A) and saccule (B) from an inner ear affected by MD (hematoxylin/eosin stain). Pathologically enlarged (EH) endolymph fluid spaces (red shadings) are compared to their normal size (grey shadings). Hydropic Reissner’s membrane (RM) and saccular membrane (SM) are highlighted (red lines) as compared to their normal configuration (grey lines). RM and SM anchoring points at rigid inner ear tissue structures are indicated (RM: strial, limbal; SM: anterior, posterior). (C–D′) High-magnification images demonstrating epithelial configurations in EH suggestive of cell multiplication: complex curling of the hydropic utricular epithelium (C′, red arrows) compared to normal utricular epithelium (C′, black arrow), and villous-like patterns of supernumerary cells at the hydropic cochlear base (D, red arrow) compared to its normally flat mono-layered epithelium (D’, black arrow). (E) Contrasting models of epithelial expansion in EH: the classic pressure-based (ΔP) mechanism predicts unchanged cell numbers (n) and increased cell spacing ((XY)) due to epithelial distension, whereas our proliferation-based model predicts primarily an increase in ‘n’ over (XY). (VIII, cochlea-vestibular nerve). Scale bars: (A, B), 500 μm; (C–D’), 100 μm.

### Revisiting historical MD inner ear specimens refutes evidence of epithelial ruptures

As an initial step, we examined two archival inner ear specimens with EH from donors with a history of MD, in which “ruptures” of hydropic epithelia were originally reported over 60 years ago (3). Utilizing the originally published tissue sections (Figs. S1A, S1A′) along with adjacent serial sections retrieved from our archive (Figs. S1B, S1B′), we performed 3D reconstructions of the hydropic epithelia at the purported rupture sites (Figs.□S1C – S1D′), arrows and light blue shaded epithelium). Contrary to the original interpretations, our 3D models revealed no epithelial discontinuities. Instead, these regions exhibited extensive epithelial puckering and outfolding along the sectioning plane, appreciable only in the 3D models. Although the very faint staining quality of these historical specimens prevented ML-based cell nucleus counting, visual inspections confirmed a dense and uniform distribution of cell nuclei within the outpouchings, suggesting no pressure-induced “herniation” nor rupture-related tissue displacements, but rather structural changes driven by addition of supernumerary cells.

### Hypercellularity in hydropic inner ear epithelia is universal across disease etiologies and stages

Using machine learning (ML)-based quantification of epithelial cell nuclei (cell counts) and internuclear distances between neighboring epithelial cells (cell spacing)—an established surrogate marker for mechanical stretch acting on epithelia (26)—we found that average cell counts per cross-sectional view of RM and SM were significantly higher in both MD (idiopathic EH; RM = 169 ± 81, p < 0.0001; SM = 808 ± 386, p < 0.0001, n = 50) and secondary EH ((18); RM = 136 ± 73, p < 0.0001; SM = 556 ± 200, p < 0.0001, n = 40) compared to EH-negative controls (RM = 52 ± 13; SM = 304 ± 52, n = 100) (Figs. 2, S2). In contralateral ears (no disease history, no EH) from donors with unilateral idiopathic EH, cell counts for RM and SM were not significantly different to the EH-negative controls (RM: 51 ± 16, p > 0.05; SM: 325 ± 42, p > 0.05) (Figs. S3A, S3B). Average cell spacing in RM and SM showed no significant differences among all groups, including idiopathic EH (RM: 18.6 μm ± 14.4 μm, p > 0.05; SM: 16.7 μm ± 13.2 μm, p > 0.05), secondary EH (15.4 μm ± 11.2 μm, p > 0.05; SM: 14.2 μm ± 10.7 μm, p > 0.05), and contralateral (unaffected) ears in unilateral idiopathic EH cases (RM: 16.9 μm ± 13.6 μm, p > 0.05; SM: 13.3 μm ± 9.6 μm, p > 0.05) as compared to EH-negative controls (15.9 μm ± 11.7 μm) (Figs. 2, S2). Most specimens from the idiopathic EH group represent donors with decades-long MD in the “burnt-out” stage, characterized by moderate-to-severe permanent sensory impairment and severe end-stage EH. We separately analyzed a small group of three specimens from donors in the early, active fluctuating disease stage with milder EH (Figs. S3C–S3F). Notably, this early-stage group also showed significantly increased RM and SM cell counts (RM: 138.4 ± 49.3, p < 0.0001; SM: 831.9 ± 331.6, p < 0.01), but no significant differences in cell spacing (RM: 17.7 μm ± 12.1 μm, p > 0.05; SM: 13.6 μm ± 9.4 μm, p > 0.05) compared to EH-negative controls. This suggests that cell number increases within hydropic epithelia occur early in the dynamic phase of EH development, suggesting that hyperplasia may be a primary driver of EH formation rather than a secondary, end-stage phenomenon. Additionally, in the no EH controls, neither age nor sex correlated with changes in RM and SM cell counts or spacing, ruling out these factors as potential confounders (Figs. S3G, S3H).

**Figure 2.**
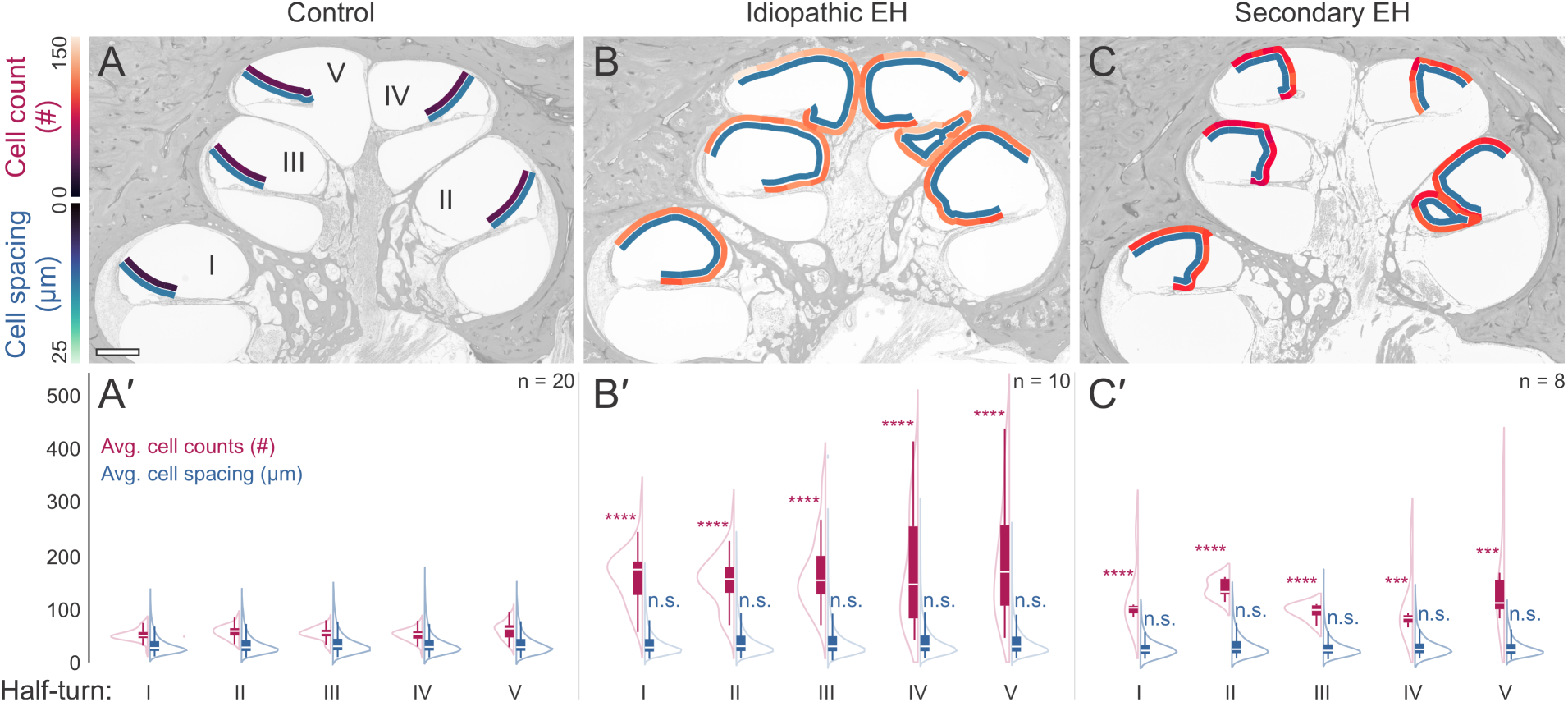
Comparison of Reissner’s membrane (RM) epithelial cellularity across EH groups. We compared RM cell counts and cell spacings in specimens from control (EH-negative), idiopathic EH, and secondary EH groups. (A, B,C) Representative heatmaps overlaying RM in each cochlear half-turn (I–V) display cell counts (top) and cell spacing (bottom) per 10% membrane segment. (A′,B′,C′) Group averages for cell counts (top) and cell spacing (bottom) are shown for comparison. See SI Appendix, Fig. S4 for heatmap generation details. **** P < 0.0001, *** P < 0.001, n.s. = not significant. Scale bar: (A–C), 500 μm.

### Focal hydrops exhibits site-restricted hyperplasia

Focal EH, confined to discrete regions along the inner ear epithelium (27), is particularly challenging to reconcile with the classic theory of fluid overpressure-induced EH formation and the underlying physical law of uniform pressure effects in closed systems (Pascal’s law, a.k.a. principle of transmission of fluid-pressure). From our archival collection of 13 rare focal EH specimens, we selected three with unequivocally severe EH: Specimens #fEH_1 and #fEH_2 both exhibited focal EH restricted to two cochlear half-turns each, while #fEH_3 showed EH solely in the saccule (Figs. 3A, 3B). In all three cases, hydropic regions showed significantly increased average epithelial cell counts compared to the corresponding areas in no EH controls, namely cochlear half-turns II (#fEH_1: 249 ± 16, p < 0.01; #fEH_2: 120 ± 11, p < 0.01) and III (#fEH_1: 187 ± 54, p < 0.05; #fEH_2: 156 ± 9, p < 0.01), and the saccule (#fEH_3: 731 ± 176, p < 0.01), while average cell spacing was not statistically different from no EH controls (half-turn II: #fEH_1: 16.1 ± 11.9 μm, p > 0.05; #fEH_2: 22.3 ± 16.6 μm, p > 0.05; half-turn III: #fEH_1: 20.1 ± 16.2 μm, p > 0.05; #fEH_2: 18.1 ± 15.4 μm, p > 0.05; saccule: #fEH_3: 15.4 ± 12.5 μm, p > 0.05). Notably, in fEH_1 and fEH_2, the non-hydropic RM in the basal cochlear half-turn (I)—immediately adjacent to the hydropic RM in half-turns II and III—exhibited significantly increased cell counts near the stria vascularis insertion point compared to control half-turn I (p < 0.05–0.001) (Figs. 3C, 3C′). These densely packed, strikingly cuboidal epithelial cells (insets, Fig. 3C) suggest that hypercellularity may precede hydropic expansion.

**Figure 3.**
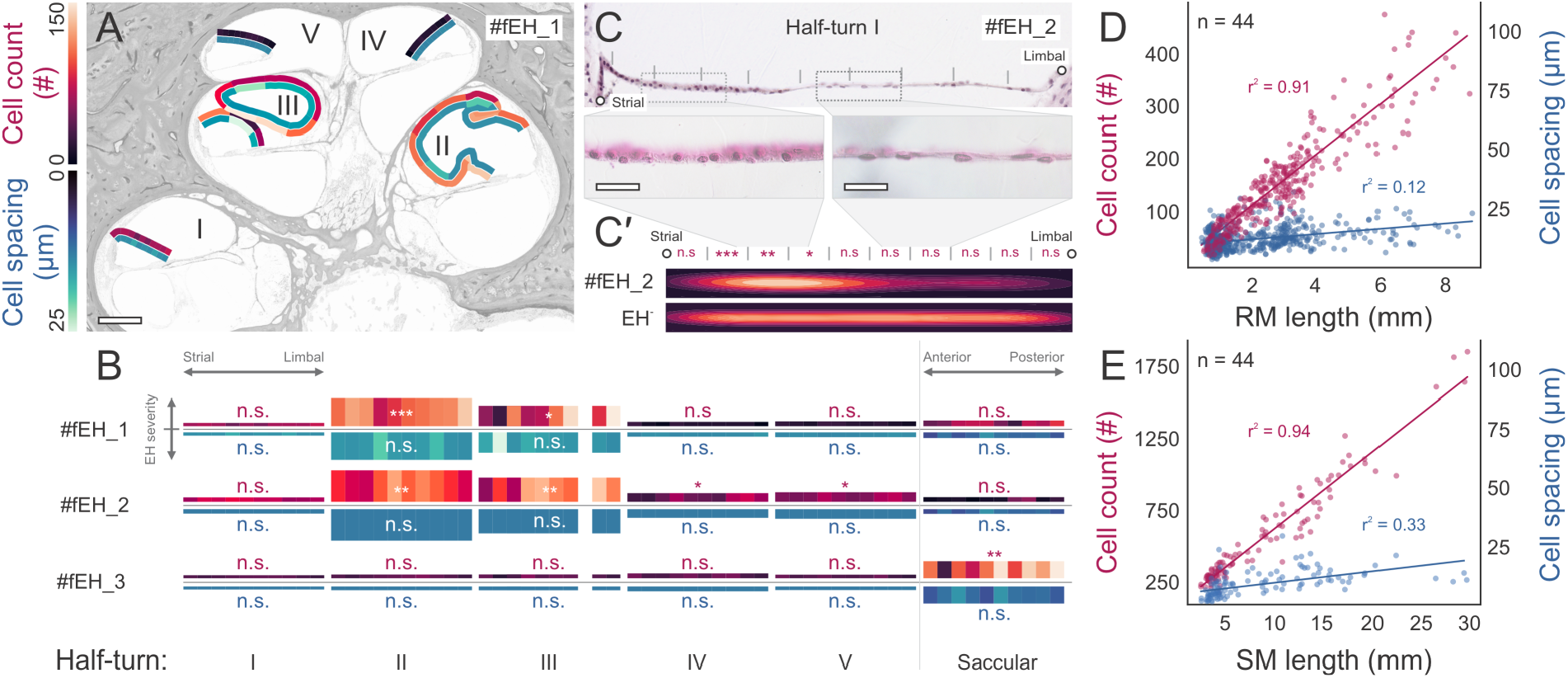
Focal endolymphatic hydrops (EH) and regional variations in epithelial cellularity. (A–C′) Reissner’s membrane (RM) and saccular membrane (SM) epithelial cell counts, cell spacing, and morphology in three focal EH (fEH_1–3) specimens. (A) Heatmaps for fEH_1 show average cell counts (top) and cell spacing (bottom) per 10% membrane segment in each cochlear half-turn (I–V). (B) Heatmaps for fEH_1–3, with heatmap height correlating with endolymph cross-sectional area (a measure of EH severity), demonstrate focal EH in cochlear half-turns II and III (fEH_1 and fEH_2) and in the saccule (fEH_3). (C, C′) High-power view of RM in cochlear half-turn I (fEH_2) reveals denser cell packing and a distinct cuboidal cell morphology near the strial end compared to the limbal end (insets). Kernel density estimate (KDE) plots of average cell counts per 10% segment show significantly higher cell counts near the stria anchoring point (see Fig. 1A for reference). (D, E) Average cell counts and cell spacing as a function of RM (D) and SM (E) length across all cases. *** P < 0.001, ** P < 0.01, * P < 0.05, n.s. = not significant. Scale bars: (A) 500 μm; (C) 20 μm.

### Cell number drives hydropic expansion of inner ear epithelia

Analysis of all specimens using linear regression (r^2^) and Pearson correlation (r) revealed a strong positive correlation between cell counts and epithelial segment lengths (RM: r^2^ = 0.90, r = 0.95; SM: r^2^ = 0.94, r = 0.97), but only a weak correlation with cell spacing (RM: r^2^ = 0.12, r = 0.35; SM: r^2^ = 0.32, r = 0.57) (Figs. 3D, 3E, S3E, S3F). These findings indicate that RM and SM cell densities remain relatively constant across all stages of hydropic expansion, supporting our hypothesis that increased cell number primarily drives hydropic expansion of endolymphatic spaces.

### Hyperplastic growth of inner ear epithelia offsets epithelial loss in diseased endolymphatic sac

To generate hypotheses about drivers of hyperplastic growth in RM and SM epithelia, we focused on the endolymphatic sac (ES)—a non-sensory inner ear region, known for developing early pathologies relevant to MD pathogenesis (23, 28). We hypothesized that these ES pathologies trigger compensatory hyperplastic mechanisms similar to tissue repair mechanisms in other organs (24, 25). To gain insight, we segmented and quantified in five healthy adult and eleven MD specimens the total endolymph-lining epithelial membrane surfaces—an established morphological correlate for epithelial (fluid and ion) transport capacity (29). In healthy inner ears, the ES accounted for 42.2% (average: 115 mm^2^, SD = 53.3 mm^2^) of the total inner ear epithelial surface (Figs. 4A, 4C). In MD specimens, the diseased ES’ epithelial surface was significantly reduced to 5.2 % (average: 15.7 mm^2^, SD = 7.7 mm^2^ ; p < 0.0001)—a 37 % loss of total inner ear epithelial surface (Figs. 4B, 4C; purple shading and data points). Conversely, other inner ear epithelia, particularly RM in the cochlea and SM in the vestibule, significantly increased through hyperplastic (hydropic) expansion by approx. 38.8 % from 157.8 mm^2^ (SD = 19.8 mm^2^) in healthy ears to 257.9 mm^2^ (SD = 45.5 mm^2^, p < 0.0001) in MD specimens (Figs. 4B, 4C; green shading and datapoints)—statistically offsetting (p > 0.05) the epithelial loss in the ES and maintaining the total combined epithelial surface of RM, SM and ES within the normal range.

**Figure 4.**
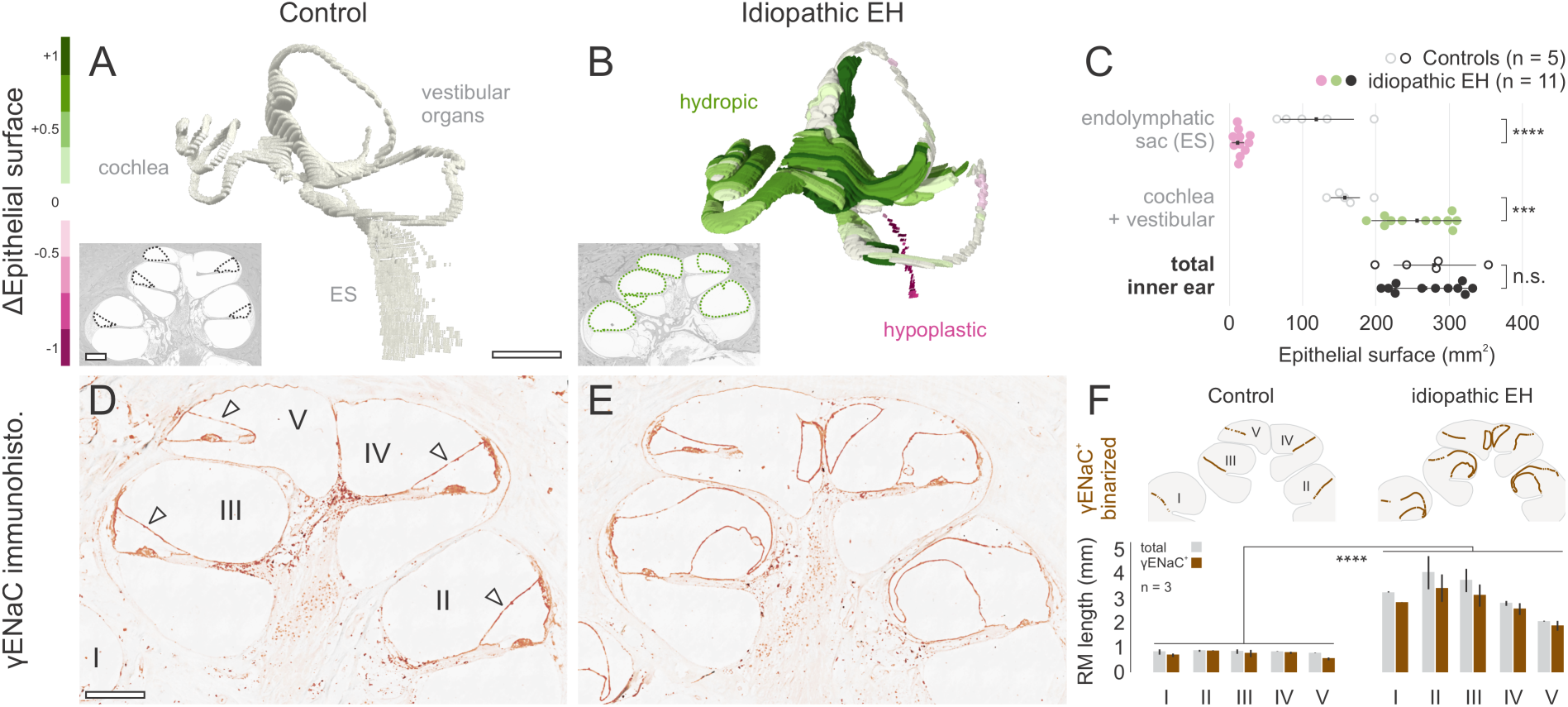
Epithelial surface area comparisons and epithelial sodium channel (γENaC) immunostaining in hydropic inner ears. (A, B) Epithelial surface comparisons between non-hydropic (A) and hydropic (B) inner ears, visualized by 2D segmentation of near-serial tissue sections (insets). The green-to-purple scalar field indicates epithelial gains (green) and deficits (purple) in the hydropic ear relative to the control. (C) Quantitative comparison of epithelial surface areas in cochleovestibular, endolymphatic sac (ES), and total (cochleovestibular + ES) portions of normal and idiopathic EH inner ears. (D, E) γENaC immunostaining in cochlear cross-sections from a non-hydropic (D) and hydropic (E) inner ear, with strongest staining in Reissner’s membrane (RM; arrowheads). Insets: γENaC localization (arrowheads) changes from apical (D, normal ES) to diminished and depolarized (E, hypoplastic ES). (F) Quantification of total and γENaC-positive RM cross-sectional lengths. Scale bars: (A, B) ≈1 cm; (insets in A, B), (D, E) 500 μm, (insets in D, E) 10 μm.

### Shared molecular features between hyperplastic inner ear epithelia and endolymphatic sac epithelia suggest functional compensation

To determine if hyperplastic growth in RM and SM generates new epithelial cells that adopt functions redundant with—and potentially compensating for—lost ES epithelial cells, we investigated shared molecular features between these epithelial sites. Specifically, we investigated the immunolocalization of the sodium channel ENaC, which has previously been reported in both rodent RM (30, 31) and ES epithelia (23, 31), where it assumes critical functions for endolymph fluid homeostasis in synergy with other ion transport proteins (32, 33). In tissue sections from a donor with MD in one ear and no disease history in the opposite ear, we found in the control ear strong ENaC immunoreactivity in both the RM and ES (Fig. 4D), contrasting with diminished ENaC in the hypoplastic ES (Fig. 4E, inset) and increased ENaC-positive area in the hydropic RM (Fig. 4E) of the diseased ear. Semiquantitative comparison of ENaC-positive RM length per cochlear half-turn demonstrated a significant increase (non-hydropic RM: 0.71 mm, SD = 0.12 mm; hydropic RM: 2.75 mm, SD = 0.58 mm; p < 0.0001) (Fig. 4F). In other words, in both non-hydropic and hydropic RMs, 88% (± 6 %, p > 0.05) of its cross-sectional length was ENaC-positive, indicating that newly added cells in hyperplastic (hydropic) RM adopt cell-type appropriate functional features—supporting the idea that hyperplastic growth indeed functionally compensates for ES pathology.

## Discussion

Our study provides novel key insights into the pathogenesis of EH and its role in MD, challenging the prevailing “endolymph overpressure and epithelial distension/rupture” paradigm. We present three major findings: (i) marked hypercellularity in hydropic epithelia without evidence of pressure-induced epithelial distension or microtrauma; (ii) a reciprocal relationship between epithelial cell loss in the ES and hyperplastic cell gain in RM and SM; and (iii) maintenance of ion transport capabilities in these newly formed cells, suggesting a compensatory rather than purely pathological process. These findings support a novel model of EH as an adaptive organ-wide response to initial ES dysfunction.

*Challenging the classic “overpressure and rupture model”*. The classic model of EH posits that an imbalance between endolymph secretion and resorption, leads to increased endolymphatic pressure, distension of RM and SM, and subsequent epithelial ruptures. These ruptures were proposed to release potassium-rich endolymph into the perilymph, disrupting mechano-electrical transduction, and synaptic transmission, thereby triggering the characteristic symptoms of MD (3). This model was largely based on historical histological reports describing such ruptures, fistulae, scarring, and herniations in inner ears affected by MD (3, 12, 34, reviewed in 35). However, physiological studies simulating rupture by perfusing animal inner ears with potassium-rich solutions (36), as well as studies of surgically inducing epithelial ruptures, failed to induce marked dysfunction of hearing or balance in these animals (37). Here, our re-evaluation of archival specimens from one of the earliest seminal reports (3) reveals that these purported rupture zones are more consistent with epithelial puckering and folding due to hyperplastic growth (Fig. S1). Furthermore, we find that tissue shrinkage and other processing artifacts, particularly prevalent in these older specimens (38), likely contributed to these misleading rupture-like appearances.

The classic model is further challenged by the phenomenon of focal EH, where hydropic changes are restricted to specific inner ear regions (e.g., distinct cochlear turns or the saccule) while adjacent areas remain unaffected (Figs. 3A, 3B). A purely hydrostatic mechanism would predict uniform pressure distribution throughout the endolymphatic system, making such focal patterns difficult to reconcile with a generalized fluid overpressure mechanism (Supplementary Discussion). Moreover, in these focal EH specimens, we observed significantly increased cellularity and abnormal cell distribution patterns in the (non-hydropic) RM of half-turns directly adjacent to hydropic areas (Fig. 3C, 3C′). This finding is critical, as it suggests that hypercellularity is a very early event, potentially preceding histologically visible EH and supporting a primary role for hyperplasia in EH development.

### A novel model: EH as a (mal)adaptive compensatory response

Our data support an alternative model: EH as an organ-wide compensatory response akin to those known from other organs like the kidney (39, 40), liver (41), and thyroid (42). In these organs, initial injury triggers hypertrophy and hyperplasia in surviving cells to preserve overall organ function. Persistent or severe injury, however, can shift this beneficial response towards a maladaptive pathway, marked by inflammation, fibrosis, and cellular senescence, ultimately leading to chronic organ failure (43). Clinically, this maladaptive transition often manifests as fluctuating or repetitively declining organ function (e.g. intermittent drops in glomerular filtration rate in early-stage chronic kidney disease), before descending irreversibly into end-stage disease (44).

Adapting these ideas to the inner ear, we propose that pathology of the ES—previously identified as an early event in MD (23)—represents the initial injury. This injury, potentially mediated by altered endolymph composition (45, 46), triggers compensatory hypertrophy and hyperplasia in RM and SM epithelia to maintain fluid homeostasis and preserve hearing and balance functions. This increased cellular activity, heightened metabolic burden, and the volume expansion of the endolymphatic fluid, which requires increased ion transport to maintain physiological ion gradients (47-49), may ultimately predispose the inner ear to degenerative changes and precipitate maladaptation. We propose that the episodic symptoms of early-stage MD reflect temporary failures of this compensatory system, similar to fluctuating/dropping function in other organs during this phase (43, 44), and its transition towards a maladaptive pathway, eventually culminating in chronic, irreversible sensorineural dysfunction of late-stage MD.

Three of our key findings support this model (i) substantial epithelial loss in the ES (reducing its contribution to the endolymph-lining epithelium from ∼42% in healthy ears to ∼5% in MD ears; Fig. 4C), comparable to renal mass loss known to trigger compensatory responses (39, 43); (ii) a reciprocal quantitative relationship between ES epithelial loss (∼37% of total inner ear epithelium) and hyperplastic expansion of RM/SM epithelia (increasing cell numbers 4-7-fold in severe EH; Figs. 3D, 3E and 4C), mirroring compensatory changes observed in other organs; and (iii) functional redundancy between lost ES epithelium and newly formed RM/SM epithelia, as evidenced by ENaC expression in a high percentage (∼88%) of both cell populations (Figs. 4D– 4F), consistent with compensatory increase of transporter proteins in other organs.

### Implications for future therapeutic research

This new model of EH as a tissue damage-driven compensatory process offers a paradigm shift for therapeutic research in MD and other EH-associated disorders. Future strategies may focus on enhancing the beneficial aspects of the compensatory response while preventing its maladaptive trajectory. Key questions for future research include: What is the precise nature of the initial inner ear pathology—in particular in the ES—that triggers EH? What specific fluid-regulatory functions do the newly added cells perform? What molecular signals govern epithelial hyperplasia, and how can these pathways be modulated to help restore homeostatic functions without exacerbating damage? Addressing these questions may lead to novel, targeted interventions that preserve inner ear function and prevent irreversible sensory decline seen in MD and related EH spectrum disorders.

## Materials and methods

### Ethics

This study was approved by the Institutional Review Board of Massachusetts Eye and Ear (IRB #2021P001593). The archival tissue specimens used were collected in compliance with ethical guidelines and with informed consent from the donors or their legal next of kin, as documented under IRB #2019P003755 and prior approvals.

### Human temporal bone tissue processing

Human temporal bones (TBs) from the Massachusetts Eye and Ear Collection were processed according to established standard methods (50). Briefly, the specimens were harvested postmortem, fixed in formalin, decalcified using EDTA, dehydrated in a graded alcohol series, and embedded in celloidin. They were then serially sectioned along a near-horizontal plane at a thickness of 20 μm, yielding approx. 500 sections per specimen. Every tenth section was stained with hematoxylin and eosin (H&E) for histological examination. The intervening unstained sections were stored in 80% ethanol for long-term preservation and use in immunohistochemical studies.

### Archival specimen selection

Based on clinical donor history and the histological severity and localization of EH, archival inner ear specimens with or without EH were assigned to six groups. (Group 1) “idiopathic EH” comprised all specimens from patients with a documented history of MD (1) and definitive histological evidence of EH. Although most specimens in this category were from patients who had advanced MD at the time of death (average disease duration 31.2 ± 19.8 (SD) years) with extensive EH, a subgroup exhibited episodic, early-disease “attack-like” symptoms at the time of death (average disease duration 8 ± 2 (SD) years) and was separately analyzed as (Group 2) “idiopathic EH_early stage.” (Group 3) “secondary EH” and (Group 4) “focal EH” encompassed specimens from patients with other clinical conditions hypothesized to induce EH (18), with EH affecting either the entire inner ear (“secondary EH”) or confined to specific locations of the endolymph-lining epithelium (“focal EH”) (27). (Group 5) “EH-negative” (control) included specimens without any histological signs of EH and no known inner ear disease; in this group, audiograms (where available) indicated age-appropriate hearing, and donors ranged in age from gestational week 28 to 104 years. Lastly, (Group 6) “idiopathic EH_contralateral” comprised specimens lacking microscopic EH taken from donors who had MD in the contralateral ear.

For the immunohistochemical analysis of epithelial sodium channel (ENaC) protein expression in hydropic and non-hydropic inner ear epithelia, we utilized the archival left and right ear specimens from the same patient: the right ear exhibited unilateral idiopathic endolymphatic hydrops (EH) and Meniere’s disease (MD), while the left ear showed no signs of EH or any clinical history of inner ear disease. This was the only recent unilateral MD case available for immunohistochemical study. Detailed clinical data for all cases are provided in Table S1.

### 3D reconstruction of alleged epithelial ruptures in historic MD specimens

Two inner ear specimens from MD patients, as described in (3), were identified in our archival specimen collection using published images from the original study. Fourteen to eighteen additional sections adjacent to those shown in the publication (both inferior and superior) were retrieved from our tissue archive. These sections were mounted on microscope slides, stained with hematoxylin and eosin (HE), and digitized along with the original stained archival slides. The digitized images were then imported into Dragonfly software (v2024.1, Comet Technologies, Montréal, Canada) as an image stack, morphologically aligned, and the endolymph-lining epithelia were manually segmented to generate 3D image representations of the areas interpreted as ruptures in the original study.

### Digital image processing

H&E-stained slides were digitized as .svs files using a Leica Aperio AT2 scanner (Leica, Wetzlar, Germany) with a 20x Plan-Apo objective at a 0.5 μm per pixel resolution. Digital images of slides showing cross-sections through all five cochlear half-turns (midmodiolar plane) and the maximum extension of the saccule were cropped in Qupath (v0.5.1, (26)) and exported as 24-bit JPEGs. For unevenly mounted (“wavy”) sections, images were scanned across 5-11 focal planes, and extended depth of field images were generated using the pyramid maximum contrast (PMax) algorithm in Zerene Stacker software (v1.04, Zerene Systems LLC, Richland, WA). For each specimen, three images from non-consecutive slides in the midmodiolar plane of the cochlea showing all five half-turns, and two to three images from slides showing the maximum extent of the saccule were selected.

### Deep-learning, semantic segmentation of cell nuclei

A U-Net semantic segmentation algorithm was trained using Dragonfly software (v 2024.1, Comet Technologies, Montréal, Canada) to infer segmentation of epithelial and mesenchymal cell nuclei in H&E-stained images of Reissner’s membrane (RM) and saccular membrane (SM) (Fig. S4A–S4M). Ground truth data were generated by manually annotating 2,342 nuclei in images from 20 archival human inner ear specimens, representing a diverse dataset processed over five decades, and encompassing variability in tissue processing artifacts and degrees of H&E staining fading anticipated in the study dataset. Training and validation datasets were augmented fivefold, stochastically modifying scaling, rotation, brightness adjustment, Gaussian noise, and flipping vertically and/or horizontally. Key hyperparameters—i.e. learning rate (0.001), learning rate decay, batch size (32), number of epochs (≤10,000), voxel-wise sampling, and the optimization algorithm (Adam)— were tailored to minimize validation loss to 0.00136 and avoid overfitting. Model performance was evaluated on a separate dataset of RM images that also represented the variability of image quality and artifacts, using as ground truth manual nuclei annotations from three independent investigators (average 1,257 +/-10 annotations per investigator), that, when averaged across the three investigators, yielded a precision of 0.96 ± 0.01, recall of 0.92 ± 0.02, F1-score of 0.94 ± 0.01, and accuracy of 0.89 ± 0.01. The trained algorithm was applied to the study dataset images that were prepared with manually drawn ROI masks of RM (in all five cochlear half-turns) and SM. To prevent overcounting in tangentially sectioned epithelial areas, adjacent endolymph spaces were segmented and dilated with a kernel size of 19 (19×19×19 voxels) to approximate cross-sectional epithelia width. Only nuclei intersecting with the dilated endolymph mask were further processed. Segmented nuclei with a 2D maximum feret diameter >10 (corresponding to flat elongated mesenchymal cell nuclei) or X/Y area pixel size <10 (dust and other image artifacts) were excluded as well. Centroid X/Y coordinates of included epithelial nuclei were exported as .csv files.

### Raw data processing

A custom Python (v3.11.10) script was used to process .csv files containing centroid X/Y coordinates of RM and SM cross-sections (Figs. S4N–S4Q). The centroids were aligned and numbered along the epithelia, from the limbal to strial end for RM and from the posterior to anterior end for SM. A nearest-neighbor heuristic was initially applied to connect neighboring centroids, forming a path that was subsequently optimized using the 2-opt algorithm to minimize overall path length; essentially ensuring that centroids were connected along the natural trajectory of the epithelium (limbal-to-strial for RM; anterior-to-posterior for SM). Two to three aligned centroid paths were generated for each RM and SM per specimen, based on two to three (n = 2-3) RM and SM images processed per specimen. The following metrics were calculated: the average total number of centroids (“average cell counts”), the average neighboring centroid distance (“average cell spacing”), and the average sum of neighboring centroid distances (“average total RM and SM lengths”). Heatmaps showing average cell counts and cell spacing per 10% path segment were generated and overlaid onto the corresponding RM and SM segments in the original microscope images.

### Comparative epithelial surface area analysis in hydropic and non-hydropic inner ears

To quantitatively compare epithelial surface area expansion due to EH and surface area loss due to endolymphatic sac pathology in inner ears with MD (idiopathic EH), we measured the two-dimensional surface areas (mm^2^) of all endolymph-lining epithelia in the inner ears of eleven specimens with idiopathic EH, alongside five specimens without EH from donors of similar age without a history of otological disease. The epithelia-endolymph interfaces were manually traced on digitized sections (20□μm thickness) spaced uniformly at 200□μm intervals using QuPath (v. 0.5.1, (51)), encompassing the entire inner ear except for a small portion of the superior semicircular canal above the crista, which is routinely removed during specimen processing. These traced interfaces were then analyzed to determine surface area changes. Three-dimensional heatmap representations of the cross-sectional perimeters of the endolymph-lining epithelia in EH-affected inner ears were generated and normalized to non-EH controls. The heatmaps employed a diverging color scheme to highlight regions of epithelial surface area increase (green) and decrease (purple) for each segment.

### Immunohistochemistry

For immunohistochemical analysis on cochlear sections from a donor diagnosed with unilateral MD, tissue sections along the long-center axis of the cochlea (midmodiolar) from both the unaffected and affected inner ears were mounted side-by-side on Superfrost Plus slides and processed in parallel. The sections were de-celloidinized, rehydrated, and underwent heat-induced antigen retrieval following established protocols. Sections were blocked with 5% normal horse serum (NHS) in PBS, followed by overnight incubation with a primary antibody against the gamma-subunit of the epithelial sodium channel (γENaC) raised in rabbit (provided by J. Loffing, University of Zurich, Switzerland) and whose specificity was previously validated (23). Subsequently, a secondary biotinylated anti-rabbit IgG antibody (raised in goat, Jackson Immunoresearch, West Grove, PA) was applied for one hour at room temperature. Negative control experiments were performed by omitting the primary antibody, which showed no staining (data not shown). The sections were then incubated synchronously in diaminobenzidine (DAB)/hydrogen peroxide in PBS, supplemented with 4% DAB (Sigma), for two to ten minutes. Finally, the slides were dehydrated through a graded ethanol series, cleared with Histo-clear (National Diagnostics, Atlanta, GA), and coverslipped using Permount (Fisher Scientific, Waltham, MA).

### Microscopic analysis

HE-stained tissue and DAB-labeled tissue sections were examined using differential interference contrast on a Nikon Eclipse E800 microscope (Nikon, Tokyo, Japan). Images for comparing ENaC-positive RM lengths were captured with consistent microscope settings, including a 10x objective, fixed exposure time, and camera gain.

### Quantification of γENaC-immunopositive epithelial segment lengths

Microscope images of DAB-labeled RM segments in all five cochlear half-turns were analyzed using Fiji/ImageJ2 (v2.14.0/1.54f, (52)). Each image was binarized by setting the black-white threshold at the pixel intensity level of γENaC-negative (DAB-negative) regions within the same images. Following binarization, histograms were generated along manual tracings of the RM lines to quantify the lengths of black (γENaC-positive) and white (γENaC-negative) pixels. The cumulative lengths of both γENaC-positive and γENaC-negative pixels were calculated for each RM segment. This analysis was performed on three non-consecutive tissue sections from each of the two inner ear specimens, and the data were pooled for subsequent statistical analysis.

### Statistics

Normality of all datasets was assessed using the Shapiro-Wilk test. Independent two-sample t-tests were used to compare average cell counts and cell spacing per cross-sectional RM and SM segments between groups with EH (idiopathic, secondary, focal, idiopathic-active) with no EH (controls) and idiopathic-contralateral respectively, manually segmented epithelial surface areas of entire inner ears with idiopathic EH and no EH (controls), and γENaC-positive immunoreactivity lengths in RM and SM cross-sections between idiopathic EH and controls. Statistical significance was set at p < 0.05, with levels of significance indicated by asterisks: p < 0.05 (*), p < 0.01 (**), p < 0.001 (***), and p < 0.0001 (****). All analyses were conducted using GraphPad Prism (v10.3.1, GraphPad Software, San Diego, CA).

## Supporting information

Supplementary discussion

## Acknowledgments and funding sources

We gratefully acknowledge Diane Jones, Barbara Burgess, and Clarinda Northrop for their decades of dedicated work in the Otopathology Laboratory, particularly in preparing the human temporal bone specimens essential to this study. We thank J.C. Adams for intellectual contributions to the project’s early development, M.C. Liberman for valuable manuscript input, and Nanmaran Anbuselvan for technical assistance with segmentation and 3D reconstruction. This study was supported by National Institute on Deafness and Other Communication Disorders (U24-DC020849, U24-DC013983) and the Zürcher Stiftung für das Hören (Zürich, Switzerland).

D.B. was supported by a national MD-PhD scholarship from the Swiss National Science Foundation (SNSF, #183978).

