## Supplementary material for "Hyperplastic Growth, Not Hydrostatic Distension, in Endolymphatic Hydrops in Humans Challenges the Classic View of Meniere’s Disease": Figure S1

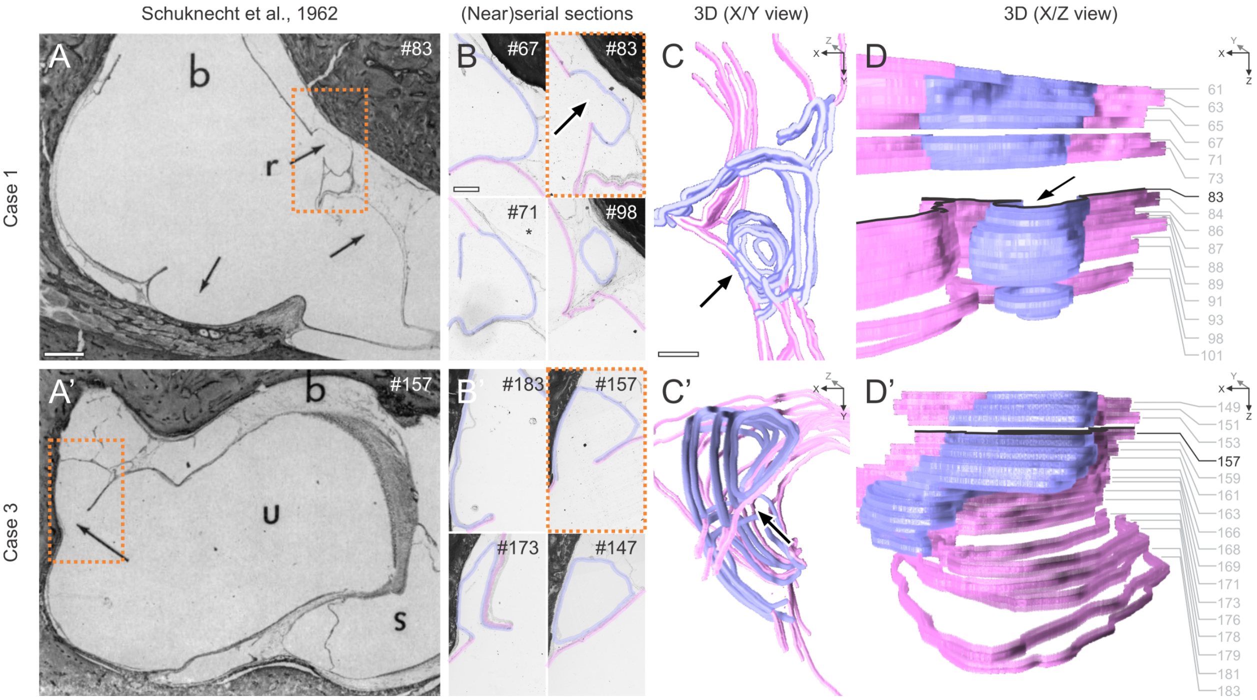


**Fig. S1.** Reinterpretation of previously reported "ruptures" in hydropic epithelia. (A, A′) Original images from (11); reproduced with permission from SAGE Publishing) showing areas within hydropic epithelia previously interpreted as ruptures (arrows) in two MD patient temporal bones. (B, B′) Adjacent tissue sections from these archival specimens (64 and 67 years old) were retrieved, mounted, digitized, and segmented. Alleged rupture sites (light blue) and neighboring epithelial regions (pink) were reconstructed in 3D (680 µm and 800 µm depth, respectively). (C, C′, D, D′) X/Y projections (C, C′; arrows indicate alleged ruptures) and Z/Y projections (D, D′) reveal no evidence of epithelial discontinuity but rather puckering and outfolding. Scale bars: (A, A′), 500 µm; (B, B′), (C, C′), 200 µm.
