## Supplementary material for "Hyperplastic Growth, Not Hydrostatic Distension, in Endolymphatic Hydrops in Humans Challenges the Classic View of Meniere’s Disease": Figure S2

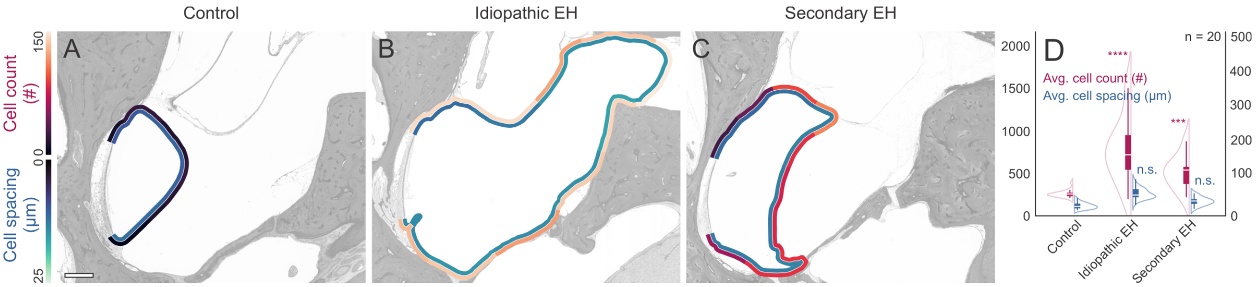


**Figure S2.** Saccular membrane (SM) epithelial cell counts and cell spacing were quantified in specimens from control (EH-negative), idiopathic EH, and secondary EH groups. (A–C) Heatmaps overlaying SM show cell counts (outer heatmaps) and cell spacing (inner heatmaps) per 10% membrane segment. (D) Group averages for cell counts and cell spacing. **** P < 0.0001, *** P < 0.001, n.s. = not significant. Scale bar, 500 μm.
