## Supplementary material for "Hyperplastic Growth, Not Hydrostatic Distension, in Endolymphatic Hydrops in Humans Challenges the Classic View of Meniere’s Disease": Figure S3

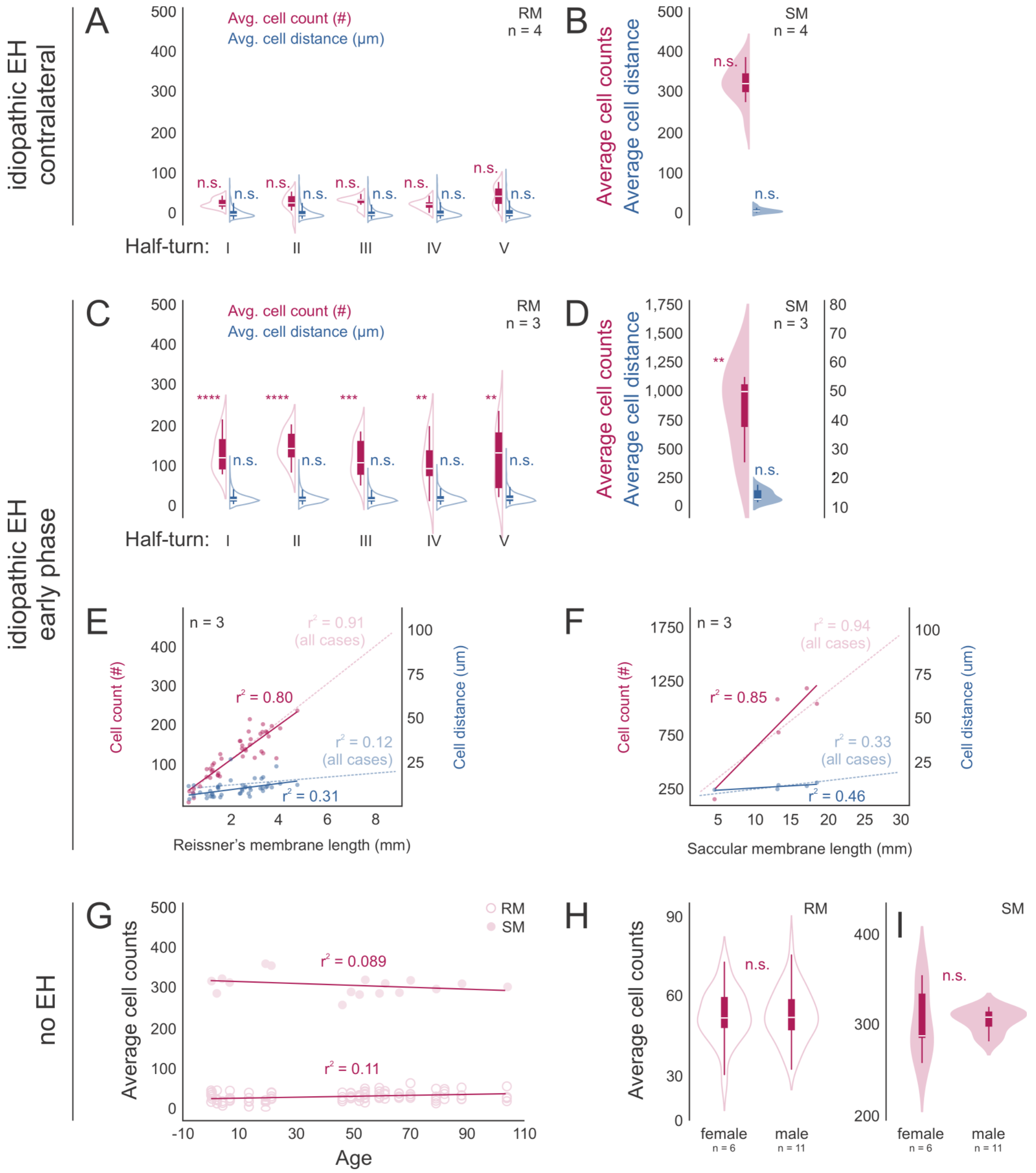


**Fig. S3.** Cellularity in contralateral ears of MD patients, early-stage idiopathic EH, and controls (EH-negative). (A, B) In donors with unilateral idiopathic EH and a clinical history of MD, contralateral (no MD, no EH) ears show RM (A) and SM (B) cell counts and cell spacing in each cochlear half-turn that are not significantly different from the no EH control group (Fig. 2A′, SI Appendix, Fig. S3D). (C, D) Early-stage idiopathic EH (clinical MD) donors, exhibiting episodic symptoms and milder EH, show RM (C) and SM (D) cell counts and cell spacing. (E, F) Cell counts and cell spacing are plotted against membrane length in RM (E) and SM (F), with the entire idiopathic EH cohort (all cases combined) shown for comparison. (G) In individuals without EH or documented inner ear disease (controls), correlations of average RM/SM cell counts with donor age (gestational week 28 to 104 years) are shown. (H, I) Distribution of average cell counts as a function of sex. **** P < 0.0001, *** P < 0.001, ** P < 0.01, n.s. = not significant.
