## Supplementary material for "Hyperplastic Growth, Not Hydrostatic Distension, in Endolymphatic Hydrops in Humans Challenges the Classic View of Meniere’s Disease": Figure S4

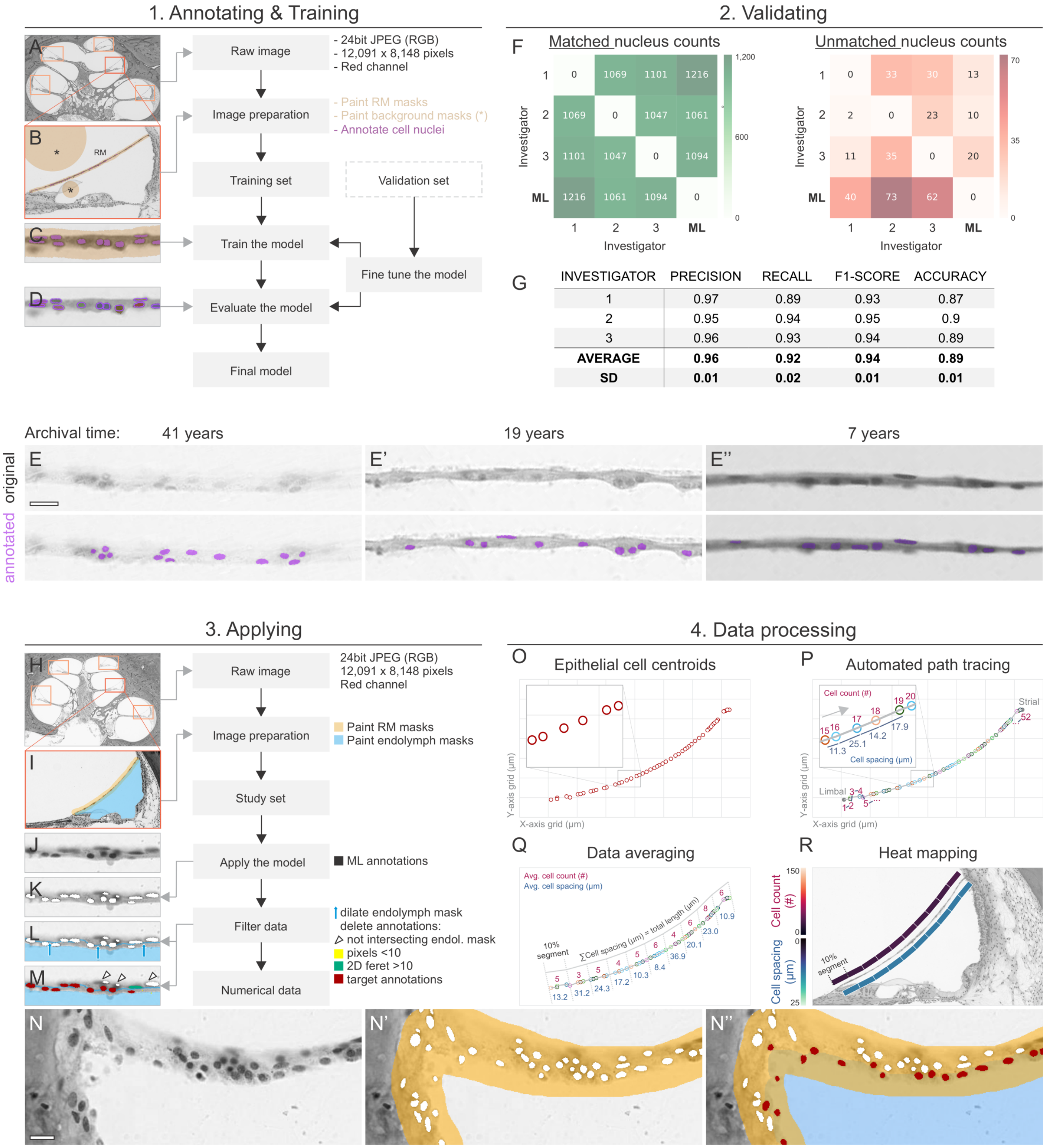


| INVESTIGATOR | PRECISION | RECALL | F1-SCORE | ACCURACY |
|---|---|---|---|---|
| 1 | 0.97 | 0.89 | 0.93 | 0.87 |
| 2 | 0.95 | 0.94 | 0.95 | 0.9 |
| 3 | 0.96 | 0.93 | 0.94 | 0.89 |
| **AVERAGE** | **0.96** | **0.92** | **0.94** | **0.89** |
| **SD** | **0.01** | **0.02** | **0.01** | **0.01** |

**Fig. S4.** Machine-learning (ML) model training, validation, and data analysis workflow. (A– E″) To train our U-Net-based ML model for segmenting epithelial cell nuclei in Reissner's membrane (RM) and the saccular membrane (SM), we manually annotated nuclei in both membranes from a range of archival human inner ear specimens covering over 40 years of storage. This broad timespan led to variability in H&E staining intensity and tissue artifacts (e.g., yellowing mounting media, fading HE staining). Hence, we curated two separate training datasets—one comprising more recent specimens and the other encompassing older, more artifact-prone specimens—to ensure robust model performance (E–E″). (F–G) Model validation was carried out on an independent heterogeneous dataset of RMs. Three investigators manually labeled >1,200 nuclei, serving as ground truth. A custom Python script employed a heuristic nearest-neighbor approach to match centroid coordinates between the ML-based segmentation and the investigator annotations, generating confusion matrices for matched and unmatched nucleus counts (F). The model yielded high precision, recall, F1 scores, and accuracy when compared to each investigator, reflecting a close match to human expert segmentation (G). (H– N″) After training, the ML model was applied to each cross-sectioned RM (H) and SM (not shown) in the study dataset. Each membrane outline (RM outlined yellow, I) and its adjacent endolymph space (blue, I) were traced. To count only epithelial nuclei directly bordering to the endolymph space, the latter was “dilated” (kernel size of 19 px; K, blue arrows). Dust or minute artifacts were removed if they had fewer than 10 pixels (yellow in M), and elongated mesenchymal nuclei were excluded if they exceeded a Feret diameter of 10 μm (green in M). This filtration in particular helped avoid double-counting multiple epithelial nuclear rows in tangentially sectioned membranes (especially in hydropic areas), confining the analysis to a single row of epithelial nuclei (N–N″). (O–R) We exported the centroid (X/Y) coordinates of counted epithelial nuclei as .csv files for subsequent analysis (O). A custom Python script identified the start and end centroids (e.g., limbal to strial edges for RM) and reordered the coordinates into a path (P). Each RM or SM segment was then subdivided into 10 equal subsegments, and for each subsegment we calculated the average number of cell nuclei (cell counts) and their average cell spacing (Q). These values, averaged over three microscope images per specimen and over entire study groups, were visualized as heatmaps that were overlaid on the original microscope images to show spatial patterns of cell density (R). Scale bars: (E–E″, N–N″), 20 μm.
