## Supplementary material for "Hyperplastic Growth, Not Hydrostatic Distension, in Endolymphatic Hydrops in Humans Challenges the Classic View of Meniere’s Disease": Table S1

Table S1. Medical record data associated with human temporal bone specimens.

| Study group | Case # | Age  (years) | Sex | Otological diagnosis | Time from diagnosis to death (years) | Specimen side | ES pathology |
| --- | --- | --- | --- | --- | --- | --- | --- |
| EH-negative (control) | No EH_1 | 28 * | Unknown | None | N.a. |  | None |
|  | No EH_2 | 0.003 | M | None | N.a. | R | None |
|  | No EH_3 | 1.83 | M | None | N.a. | R | none |
|  | No EH_4 | 3.75 | M | None | N.a. | L | None |
|  | No EH_5 | 6.37 | M | None | N.a. | R | None |
|  | No EH_6 | 13 | F | None | N.a. | L | None |
|  | No EH_7 | 19 | F | None | N.a. | R | None |
|  | No EH_8 | 21 | F | None | N.a. | R | None |
|  | No EH_9 | 46 | F | None | N.a. | R | None |
|  | No EH_10 | 49 | F | None | N.a. | L | None |
|  | No EH_11 | 52 | M | None | N.a. | R | None |
|  | No EH_12 | 54 | M | None | N.a. | L | None |
|  | No EH_13 | 59 | F | None | N.a. | R | None |
|  | No EH_14 | 61 | M | None | N.a. | R | None |
|  | No EH_15 | 66 | F | None | N.a. | L | None |
|  | No EH_16 | 70 | M | None | N.a. | L | None |
|  | No EH_17 | 79 | M | None | N.a. | L | None |
|  | No EH_18 | 82 | F | None | N.a. | L | None |
|  | No EH_19 | 88 | M | None | N.a. | R | None |
|  | No EH_20 | 104 |  | None | N.a. |  | None |
| Idiopathic EH | IEH_1 | 58 | M | Bilateral MD | 26 (R, L) | R + L | Degeneration |
|  | IEH_2 | 65 | F | Bilateral MD | 6 (R, L) | R + L | Hypoplasia |
|  | IEH_3 | 71 | M | Right MD | 11 | R | Degeneration |
|  | IEH_4 | 81 | M | Bilateral MD | 24 (L)  26 (R) | R + L | Hypoplasia |
|  | IEH_5 | 85 | M | Right MD | 20.5 | R | Hypoplasia |
|  | IEH_6 | 87 | F | Right MD | 15 | R | Hypoplasia |
|  | IEH_7 | 90 | M | Left MD | 38 | L | Hypoplasia |
|  | IEH_8 | 91 | F | Bilateral MD | 51 (R, L) | R + L | Hypoplasia |
|  | IEH_9 | 94 | M | Bilateral MD | 30 (R, L) | L | Hypoplasia |
|  | IEH_10 | 96 | M | Bilateral MD | 69 (R, L) | R + L | Hypoplasia |
| Idiopathic EH_early stage | IEH_11_early stage | 15 | F | Right MD | 10 | R | Degeneration |
|  | IEH_12_early stage | 58 | M | Right MD | 8 | R | Hypoplasia |
|  | IEH_13_early stage | 68 | F | Left MD | 6 | L | Hypoplasia |
| Idiopathic EH_contralat. | IEH_3.2_contralat. | 71 | M | Right MD | N.a. | L | None |
|  | IEH_5.2_contralat. | 85 | M | Right MD | N.a. | L | None |
|  | IEH_6.2_contralat. | 87 | F | Right MD | N.a. | L | None |
|  | IEH_7.2_contralat. | 90 | M | Left MD | N.a. | R | None |
| Secondary EH | SEH_1 | 23 | F | Labyrinthitis | N.a. | R | None |
|  | SEH_2 | 40 | M | Von Hippel-Lindau (ES tumor) | N.a. | L | None |
|  | SEH_3 | 56 | M | Cholesteatoma with erosive labyrinth fistula, post-surgical | N.a. | R | None |
|  | SEH_4 | 62 | F | Cogan’s syndrome | N.a. | R | None |
|  | SEH_5 | 63 | M | AIED | N.a. | L | None |
|  | SEH_6 | 69 | M | Idiopathic SNHL | N.a. | R | None |
|  | SEH_7 | 74 | F | Otosclerosis | N.a. | L | None |
|  | SEH_8 | 75 | M | Neurofibromatosis type 2 (schwannoma VIII) | N.a. | R | None |
| Focal EH | fEH_1 | 76 | F | Idiopathic SNHL | N.a. | L | None |
|  | fEH_2 | 78 | M | Idiopathic SNHL | N.a. | L | None |
|  | fEH_3 | 91 | F | Idiopathic SNHL | N.a. | R | None |

(ES, endolymphatic sac; SNHL, sensorineural hearing loss; AIED, autoimmune inner ear disease; MD, Meniere’s disease; R, right specimen; L, left specimen; n.a., not applicable).
