## Supplementary discussion for "Hyperplastic Growth, Not Hydrostatic Distension, in Endolymphatic Hydrops in Humans Challenges the Classic View of Meniere’s Disease"

Supporting discussion

**Origins of hypercellularity in hydropic epithelia.** The 4- to 7-fold increase in epithelial cell counts we observed in hydropic Reissner’s membrane (RM) and saccular membrane (SM) strongly suggests that intrinsically increased epithelial proliferation (hyperplasia) is the primary source of these additional cells. Ideally, this would be validated by e.g. demonstrating an increase in mitotic figures in hydropic epithelia compared to controls. However, several biological and methodological limitations inherent to our archival postmortem tissue specimens precluded such a mitotic index analysis. First, EH expansion likely follows saturating kinetics, meaning that specimens from late-stage disease—the stage represented by most of our archival material—exhibit approximately the same degree of maximal EH expansion. Consequently, the proliferation index in hydropic epithelia, presumably already low in the early, dynamic phase of EH development, has likely declined further, or even ceased entirely, by this late stage, making the presence of mitotic figures a relatively unlikely event. Second, the fact that our archival collection contains only every 10^th^ tissue section (meaning only 10% of the total epithelium could be examined), combined with the historical choice of cross-sectional angles optimized to visualize the flat epithelium en face, further limits the chance of detecting mitotic figures.

While less likely, alternative biological mechanisms underlying the increased cell counts in hydropic epithelia include mesenchymal-epithelial transition (MET, (1)), where mesenchymal cells from the perilymph-facing surface of the RM and SM would adopt epithelial characteristics. However, the baseline mesenchymal cell numbers we quantified in RM and SM from control specimens (data not shown) are insufficient by a factor of 10^2^ (data not shown) to account for the observed 4- to 7-fold cell increase in hydropic epithelia. Another possibility is bone marrow-derived cells or other cells migrating and transdifferentiating into epithelial cells (2). However, such cell migration patterns should be readily apparent given the exposed, free-floating configuration of RM and SM within fluid spaces, but was not observed by us, particularly not in early-stage EH. Therefore, while other possibilities cannot be fully excluded, intrinsic epithelial proliferation (hyperplasia) remains the most plausible explanation for the observed hypercellularity. Definitive determination of the underlying mechanism will require in vivo animal studies employing lineage tracing and proliferation markers to distinguish between proliferation and other potential processes.

**Cell spacing in hydropic epithelia.** The weak but significant correlation between cell spacing (internuclear distance) and epithelial length we observed in both Reissner's membrane (RM) (r² = 0.12, r = 0.35) and the saccular membrane (SM) (r² = 0.32, r = 0.57) (Figs. 3D, 3E) presents some interpretive uncertainty. Most likely this result is attributable to a methodological artifact. Specifically, with increasing EH severity, the expanding RM and SM epithelia come into closer proximity, and eventually abut, the surrounding tissues. In our digital image analysis workflow, this phenomenon ultimately resulted in increased pixel information noise (cell nuclei more challenging to delineate for machine-learning algorithm from surrounding tissue components), occasionally leading to missed nuclei, and artificially inflating cell spacing measurements. Consistent with this hypothesis, increase in cell spacing was most evident in the most severe EH cases upon visual inspection by the investigators. Other potentially relevant biological factors to consider that could lead to an increase in cell spacing are fluid pressure—via passive mechanical cell distension as proposed by the old model—and hypertrophy—via active increase in cytoplasm volume and membrane surface. While our findings challenge the concept of pressure as a primary pathological driver of EH, specifically refuting previously alleged pressure-induced epithelial ruptures, pressure may nonetheless play a role as an “intermediary”. In our model, disturbed fluid homeostasis (resulting from osmotic and fluid volume shifts due to endolymphatic sac dysfunction) leads to pressure changes that transduce this imbalance into a mechanical stimulus. This stimulus, in turn, may trigger the compensatory response, analogous to the effects of increased blood load in the heart (3) and urine load in the kidney (4). Hypertrophy (increased cell size, membrane surface area, and gene expression contributing to functional compensation (5)), often synergistic with hyperplasia, represents another potential contributor to increased cell spacing. Our model explicitly considers for the first time hypertrophy as a possible phenomenon in hydropic inner ear epithelia. Hypertrophy, via increase in cytoplasm volume and membrane surface are, could potentially show up as elongated cells, leading to the same morphological changes (increased cell spacing) traditionally attributed to pressure-induced mechanical distension. However, inherent limitations in our use of postmortem tissue precluded definitive demonstration of hypertrophy in this study. Consequently, histological analysis alone cannot distinguish between passive pressure effects and active cell hypertrophy.

**Mechanisms and sites of primary inner ear injury in secondary EH.** Secondary EH is typically diagnosed histopathologically when inner ear symptoms (not necessarily MD symptoms) and EH are observed alongside other conditions that affect the inner ear, including labyrinthitis, autoimmune diseases, and temporal bone trauma, and others (6). While some of these conditions directly impact the ES—for example, temporal bone fractures disrupting the vestibular aqueduct (7), focal fibrous dysplasia (8), or meningiomas (9) destroying or compressing the ES—others likely exert their pathomechanistic effects at different inner ear sites, potentially involving diffuse or even sub-light microscopic damage. Consequently, the inner ear, including the ES, may even appear histologically unremarkable in some instances. Within our novel model of EH as a compensatory hyperplastic response to inner ear injury, we propose—analogous to chronic kidney disease, which can arise from a host of insults (inflammatory, traumatic, toxic, genetic)—that various agents can damage different and/or multiple cellular sites critical for inner ear homeostasis, all leading to the same (histologically) observable compensatory response: hyperplastic EH formation. Whether this response is also consistent at the molecular level requires further investigation. This model not only redefines EH as an active participant in the pathogenesis of these inner ear conditions, moving it beyond its previous label as a mere epiphenomenon (10), but also identifies EH as a unifying response to major injury of inner ear homeostatic functions, suggesting a common pathogenic mechanism shared by MD and other seemingly disparate inner ear conditions.

**Study limitations.** Our analysis of archival, postmortem human inner ear specimens, representing a single end-stage disease time point, is retrospective, precluding direct causal inferences and assessments of dynamic relationships between hyperplasia and EH development. However, the observation of epithelial hyperplasia in both early-stage idiopathic EH (MD) specimens and non-hydropic regions adjacent to focal EH strongly suggests that hyperplasia is an early, potentially driving, factor in EH development, rather than a secondary reaction. This warrants further investigation in animal models with inducible EH to study the earliest and dynamic phases of EH development. Furthermore, our immunohistochemical analysis of ENaC was limited to two paired specimens from a single donor with unilateral MD due to decades-long archival times and variability in tissue processing, which compromised antigen preservation. Future studies employing transcriptomic analyses of postmortem human inner ears from MD patients and, crucially, functional studies in EH animal models are necessary to more comprehensively elucidate whether hydropic/hyperplastic changes are consistent with our proposed compensatory response of inner ear epithelia.
